# Cardiac glycosides cause selective cytotoxicity in human macrophages and ameliorate white adipose tissue homeostasis

**DOI:** 10.1101/2020.09.18.293415

**Authors:** Antoni Olona, Charlotte Hateley, Ana Guerrero, Jeong-Hun Ko, Michael R Johnson, David Thomas, Jesus Gil, Jacques Behmoaras

## Abstract

Cardiac glycosides (CGs) inhibit the Na^+^,K^+^-ATPase and are widely prescribed medicines for chronic heart failure and cardiac arrhythmias. Recently, CGs have been described to induce inflammasome activation in human macrophages, suggesting a cytotoxicity that remains to be elucidated in tissues. Here we show that human monocyte-derived macrophages (hMDMs) undergo cell death following incubation with nanomolar concentrations of CGs, and in particular with ouabain (IC_50_=50 nM). The ouabain-induced cell death is more efficient in hMDMs compared to non-adherent PBMC populations and is through on-target inhibition of Na,K-ATPAse activity, as it causes an intracellular depletion of K^+^, while inducing accumulation of Na^+^ and Ca^2+^ levels. Consistently, the cell-death caused by these ion imbalances can be rescued by addition of potassium chloride in hMDMs. Strikingly, white adipose tissue (WAT) explants from morbidly obese patients cultured with nanomolar concentrations of ouabain causes depletion of macrophages, decreases type VI collagen levels, and ameliorates insulin-sensitivity *ex vivo*. These results suggest that the usage of nanomolar concentration of CGs can be an attractive therapeutic avenue in metabolic syndrome characterised by pathogenic infiltration and activation of macrophages.

## Introduction

The sodium-potassium pump (Na^+^,K^+^-ATPase) is responsible for establishing Na^+^ and K^+^ concentration gradients across the plasma membrane in mammalian cells. This is the largest protein complex of the P-type family of cation pumps which hydrolyses ATP to allow the transport of K^+^ ions inside and Na^+^ ions outside cells in a 2:3 stoichiometry (Kaplan, 2002). Cardiac glycosides (CGs) are potent and highly specific inhibitors of Na^+^,K^+^-ATPase. They form a large family of naturally derived organic compounds presenting sugar and steroid moieties and have been used to treat heart conditions for more than two centuries (Prassas and Diamandis, 2008). Although toxic at a certain threshold, CGs exert positive inotropic effects, which translate into an increase of intracellular calcium ions – essential for improving the contractile function of the heart. Calcium ions are also crucial regulators of cell death (Zhivotovsky and Orrenius, 2011), which makes CGs cytotoxic.

CG cytotoxicity causes different forms of cell death (apoptosis or necrosis) in cells other than cardiomyocytes (Xiao et al., 2002) and in particular in metastatic cells (McConkey et al., 2000). Thus CGs have been proposed as an alternative therapy in cancer through immunogenic cell death (Menger et al., 2012) and apoptosis of senescent cells (Guerrero et al., 2019; Triana-Martinez et al., 2019).

One particular cell type that is particularly sensitive to intracellular potassium levels is the macrophage. Low concentration of potassium activates the Nlrp3 inflammasome and causes the maturation and secretion of IL-1β (Petrilli et al., 2007). The decrease in intracellular K^+^ can be induced by ATP and other damage-associated molecular patterns (Franchi et al., 2007; Munoz-Planillo et al., 2013; Petrilli et al., 2007) and K^+^ efflux is through membrane channels such as P2RX7 (Chen and Nunez, 2010) or TWIK2 (Di et al., 2018). Activation of P2×7 receptor (P2RX7) also induces TNF release in macrophages (Barbera-Cremades et al., 2017), suggesting that consequences of a decrease in intracellular K^+^ can be broader than Nlrp3-mediated IL-1β production.

Although CGs also decrease the intracellular K^+^, their exact role in macrophage activation and death is more controversial, presumably because of the distinct sensitivity of murine Na^+^,K^+^-ATPase to CGs (Perne et al., 2009; Price and Lingrel, 1988) and the potential implication of Na^+,^K^+^-ATPase in signalling events independently of ion transport (Cavalcante-Silva et al., 2017). In human PBMCs, monocytes and macrophages, nanomolar concentrations of ouabain stimulates pro-inflammatory cytokine expression, including IL-1β and TNF through NF-κB (Chen et al., 2017; Foey et al., 1997; Matsumori et al., 1997; Teixeira and Rumjanek, 2014) but only recent reports linked CG-induced cytokine release to inflammasome-mediated cell death (i.e. pyroptosis) (Kobayashi et al., 2017; LaRock et al., 2019).

Here we show that human monocyte-derived macrophages (hMDMs) undergo cell death following incubation with nanomolar concentrations of CGs, and in particular with ouabain (IC_50_=50 nM). We show that non-adherent PBMCs do not show the same sensitivity to ouabain-induced cytotoxicity and the sugar moiety the molecule is essential in driving CG-mediated cell death. We also report that ouabain-induced cell death is through the canonical activity of Na^+^/K^+^ ATPAse. In a proof-of-principle study, we then used ouabain-mediated selective macrophage killing in a setting where these cells cause tissue damage and persistent inflammation and fibrosis. White adipose tissue (WAT) explants from morbidly obese patients cultured with nanomolar concentrations of ouabain caused almost complete depletion of macrophages, decreased type VI collagen levels, while ameliorating insulin-sensitivity *ex vivo*. These results suggest that the usage of nanomolar concentration of CGs can be an attractive therapeutic avenue in metabolic syndrome characterised by pathogenic infiltration and activation of macrophages in the white adipose tissue.

## Methods

### White adipose tissue (WAT) explants ex vivo

Omental white adipose tissue (WAT) samples were obtained from patients undergoing bariatric surgery in St Mary’s Hospital, London. Informed consent was obtained from participants in accordance with ethical and HRA approval (REC 19/WM/0229). WAT samples were weighed and sectioned into 100 mg explants. They were then washed twice in pre-warmed HBSS containing 1% Penicillin/Streptomycin for 10 minutes each and further cultured in DMEM (Gibco) containing 10% foetal calf serum (Labtech International) and 1% penicillin-streptomycin (Sigma) with 50 nM, 100nM, 500nM of ouabain for 48 hours. For AKT, phospho-AKTser473 and CD36 Western blots, WAT explants were cultured in adipocyte maintenance media (PromoCell) containing insulin. The media was replaced with fresh media containing insulin 2 hours before harvesting the WAT explants.

### Primary adipocyte culture from human white adipose tissue

Human omental WAT was digested in HBSS containing 0.5% BSA, 10mM CaCl_2_ and 4mg/ml collagenase type I (Sigma) in constant shaking at 37°C for 1 hour. The stromal vascular fraction (SVF) was isolated from the adipose fraction by centrifugation at 300 rpm for 10 minutes. The SVF was then washed twice in HBSS and cultured with DMEM containing 10% foetal calf serum and 1% penicillin-streptomycin. For fibroblast-like stromal cell selection, the cell culture was maintained for 4 weeks and colonies were passaged once. The stromal cells were then differentiated with adipocyte differentiation media (PromoCell) until 70%-80% of cells became lipid-laden adipocyte-like cells (15 days). SVF-derived adipocytes were treated with 50nM, 100nM, 500nM and 5μM ouabain for 24 hours followed by cell viability assay.

### Isolation of macrophages and non-adherent PBMCs

Human monocyte-derived macrophages (hMDMs) were differentiated from buffy cones from healthy donors using gradient separation (Histopaque 1077, Sigma). Following Histopaque separation, peripheral blood mononuclear cells were re-suspended in RPMI 1640 (Life Technologies), and monocytes were purified by adherence for 2?hours at 37°C, 5% CO_2_. Non-adherent PBMCs were collected and washed twice with HBSS. Non-adherent PBMCs were further cultured for 5 days in RPMI 1640 containing 2-mercaptoethanol (50 nM), HEPES (10 mM), non-essential amino acid, sodium pyruvate (0.2 mM) and 20% foetal calf serum. Flow cytometry analysis showed that cultured non-adherent PBMCs were T-cells (CD3 positive; ∼75%) and non-haematopoietic (CD45 negative; ∼25%). They were then treated with 50 nM, 100nM, 500nM and 5 μM ouabain for 24 hours before performing the cell viability assay.

Adherent PBMC monolayer was washed twice with HBSS and monocytes were differentiated into hMDMs for 5 days in RPMI 1640 containing 100 ng/mL macrophage colony-stimulating factor (M-CSF, PeproTech) and 10% foetal calf serum. For the cardiac glycosides (CGs) treatment, hMDMs were treated with solvent only (Control, 0.01% DMSO), 50 nM, 100nM, 500nM and 5 μM ouabain octahydrate (Sigma), digoxin, bufalin, digitoxin and ouabagenin (Sigma), in full culture media for 24 hours. For ouabain and potassium chloride (KCl, Sigma) treatment, hMDMs were treated with KCl (25mM) 1 hour before adding ouabain (50 nM) for 24 hours. For the caspase inhibition, hMDMs were treated with 50 nM, 100nM, 500nM and 5 μM ouabain together with Q-VD-Oph (Sigma; 10μM) for 24 hours.

### Cell viability assays

For hMDMs and non-adherent PBMCs, cell viability was measured using the Annexin V and propidium iodide (PI) staining kit (BD Biosciences) according to the manufacturer’s instructions. hMDMs were detached using non-enzymatic cell dissociation buffer (Sigma) and 5×10^4^ cells were resuspended with annexin V binding buffer. hMDMs and non-adherent PBMCs were stained with annexin V (25 μg/ml) and PI (125 ng/ml) and fluorescent cells were detected using a Fortessa X20 (BD Biosciences) flow cytometer. The results were analysed using the FlowJo (v10.6) software. Duplets were excluded by gating FSC area versus FSC height and AnnexinV^-^ PI^-^ cells were considered viable cells.

For WAT-derived adipocytes, viability was measured by adding AlamarBlue (Thermofisher) directly into the culture (10%) for 4 hours. Light absorbance was measured using Multiscan Ascent (Thermo Fisher). The percentage of reduced AlamarBlue and cell viability were quantified following manufacturer’s instructions.

### Measurement of intracellular ions

hMDM were incubated with Asante Potassium Green-2 AM (Abcam, 2 μM), CoroNa Green, AM (Invitrogen, 5 μM) and Fluo-4 AM (Invitrogen, 2 μM) probes for 30 min before fixation with 1% PFA for 4 hours. Pictures were taken using an epifluorescent Olympus BX40 microscope equipped with a digital camera Retiga 2000R. Intracellular K^+^, Na^+^ and Ca^2+^ were measured from a total of 150 cells/condition from N=3 donors by quantifying the mean grey values using *ImageJ* software.

### Western blotting

WAT explants were lysed in RIPA buffer (Thermo Fisher) supplemented with protease and phosphatase inhibitor cocktails (Thermo Fisher). Total proteins were quantified by BCA protein assay (Thermo Fisher) according to manufacturer’s instructions. A total of 50μg protein was diluted 1:1 with 2x Laemmli buffer (Bio-Rad), resolved by SDS-PAGE and transferred into PVDF membranes. These were subjected to immunoblotting with the primary antibodies against phospho-Akt-Ser473 (D9E; Cell Signaling), panAkt (C67E7; Cell Signaling) CD206 (Bio-rad), CD36 (polyclonal) ACTB (Santa Cruz) and horseradish peroxidase-labelled secondary antibodies. The probed proteins were detected using SuperSignal West Femto Chemiluminescent Substrate (Thermo Fisher). Grey densities for phospho-AKT-Ser473 were analysed with *ImageJ* software and normalized by those of the loading control, ACTB.

### qRT PCR

Total RNA was extracted from hMDM and WAT explants using the TRIzol reagent (Invitrogen) according to the manufacturer’s instructions, and iScript cDNA Synthesis Kit (Bio-Rad) was used for cDNA synthesis. Quantitative RT-PCRs were performed on a ViiA 7 Real-Time PCR System (Life Technologies), using Brilliant II SYBR Green QPCR Master Mix (Agilent), followed by ViiA 7 RUO Software for the determination of Ct values. Results were analyzed using the comparative Ct method, and target mRNA expression was normalized to *HPRT* reference gene.

### Immunohistochemistry and Immunofluorescence

For CD68 immunostaining, 5 μm thick paraffin WAT sections were dewaxed and rehydrated. Antigen retrieval with sodium citrate buffer (pH 6) was carried out prior to blocking. The sections were then incubated with primary antibodies against CD68 (Bio-Rad) for 1 hour. The sections were washed in PBS, followed by the incubation with horseradish peroxidase-labeled secondary antibody (1:10,000) for 1 h at room temperature (DAKO EnVision™+ System; Agilent Technologies (UK). Diaminobenzidine (DAB) chromogen was then added, and the slides were visualized using an Olympus BX40 microscope (Olympus, UK) equipped with a digital camera Retiga 2000R CCD (QImaging, Canada).

For immunofluorescence, 5 μm thick sections were dewaxed and rehydrated. Antigen retrieval with sodium citrate buffer (pH 6) was carried out prior to blocking. Slides were then incubated overnight with Goat Anti-Type VI Collagen (Southern Biotech). After washing, slides were incubated 1 h with Donkey Anti-Goat Alexa Fluor 488 (ab150129; Abcam). Slides were mounted using VECTASHIELD medium. Images were taken using epi-fluorescent Leica DM4B microscope and the mean grey values acquired using *ImageJ* software.

### Adipocyte mean areas

WAT explants adipocyte mean areas were measured on Sirius red stained sections, using *Image J Adiposoft* plug-in (Galarraga et al., 2012). The scale was set using scale bar and the minimum and maximum areas were set to 20 and 10000 µm^2^, respectively. Pictures taken from each condition (N=4 donors; 4x magnification) were uploaded in an automatic directory processing mode and the adipocyte mean areas were calculated from a total of 8954 adipocyte areas across all the conditions.

### ELISA

Detection of human IL-1β and TNFα (Invitrogen) in hMDM culture supernatants were performed by sandwich ELISA, using technical duplicates, following the manufacturer’s instructions. Light absorbance was measured using Multiscan Ascent (Thermo Fisher) plate reader.

### Caspase-1 and caspase 3/7 activity assays

Caspase-1 and Caspase 3/7 activities were measured using Caspase-Glo® 1 and Caspase-Glo® 3/7 assay (Promega) respectively, in hMDMs and WAT explants culture supernatants, following the manufacturer’s instructions. Luminescence was measured using an Omega plate reader.

### Statistical Analysis

Results are expressed as the mean ± SEM and were analyzed using GraphPad Prism 8 software (GraphPad, USA). Statistical analyses were performed with student’s t-test, one sample t-test, or paired one-way ANOVA followed by multiple comparison Dunnett’s analysis.

## Results

### Cardiac glycosides induce cell death in human monocyte-derived macrophages (hMDMs)

We first assessed cardiac glycosides’ (CGs) cytotoxicity using Annexin V/Propidium iodide staining followed by flow cytometry analysis in hMDMs. These cells were treated with either 0.01%DMSO (control) or CGs (ouabain, digoxin, digitoxin and bufalin) at 50 nM, 100nM, 500 nM and 5μM for 24 hours. CGs-treated hMDMs showed reduced cell viability in a dose-dependent manner at nanomolar concentrations (Figure 1A and Supplementary Figure 1A). Among the CGs tested, ouabain was the most cytotoxic one (IC_50_=50) in hMDMs (Figure1A). In order to test for the cell specificity of this cytotoxic effect, we treated the non-adherent PBMCs with the same range of concentrations of ouabain for 24 hours and found that ouabain did not exert cytotoxic effects in these cells (Supplementary Figure 1B). These results indicate that hMDMs are particularly susceptible to the cytotoxic effects of nanomolar concentrations of CGs when compared to non-adherent PBMCs.

**Figure 1.**
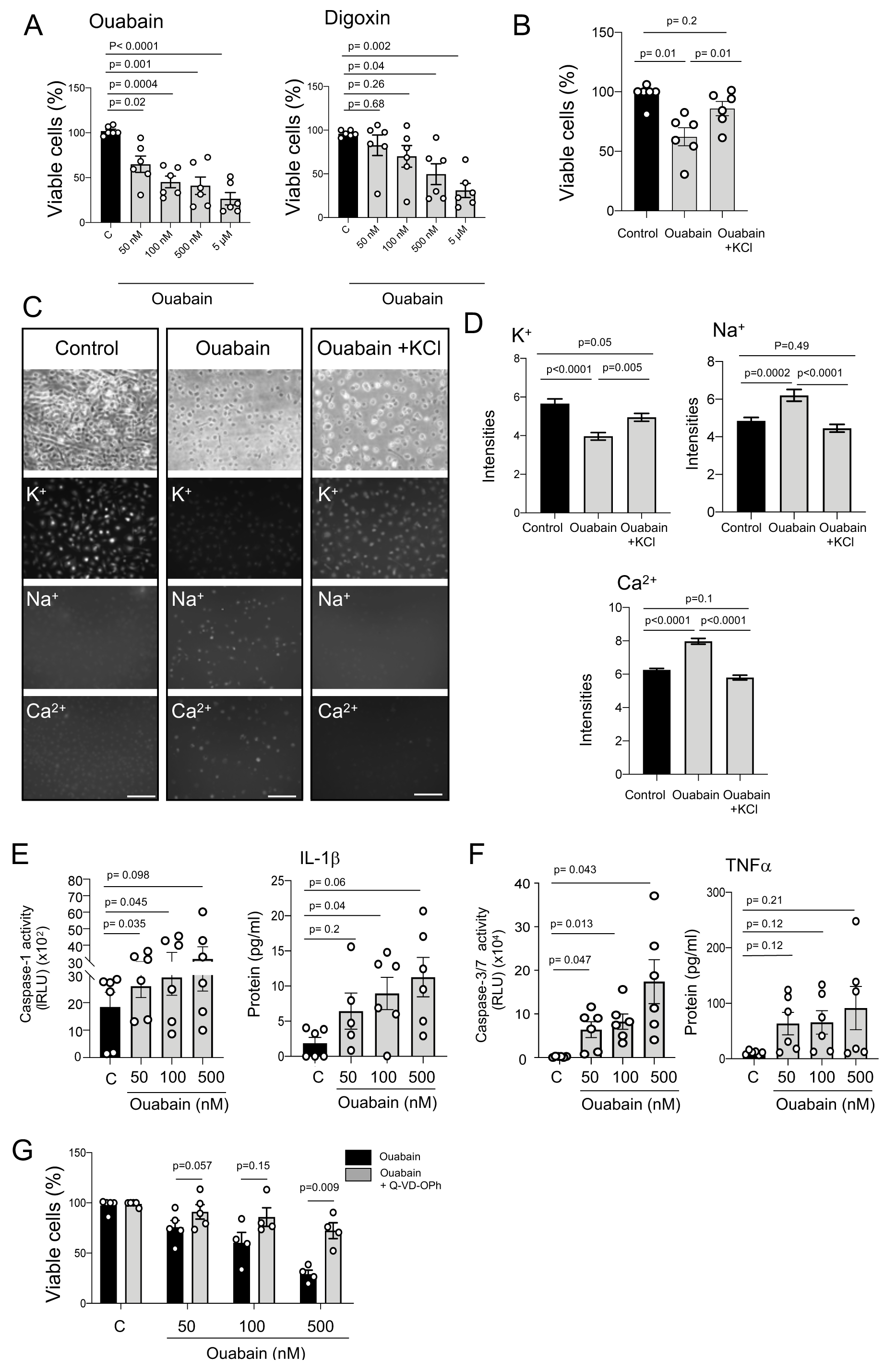
Cardiac glycosides induce caspase-dependent cell death by inhibiting the Na^+^,K^+^-ATPase in human monocyte-derived macrophages (hMDMs). (A) Cell viability (percentage of Annexin V^-^,PI^-^ cells) in hMDMs treated with ouabain and digoxin (50 nM, 100nM, 500 nM and 5 μM) for 24 hours; n=6 donors. (B) Cell viability (percentage of Annexin V^-^,PI^-^ cells) in hMDMs treated with ouabain and ouabain + KCl for 24 hours (ouabain 50 nM; KCl 25mM); n=6 donors. (C) Representative pictures of intracellular levels of K^+^, Na^+^ and Ca^2+^ in hMDMs treated with ouabain and ouabain + KCl for 24 hours (ouabain 50 nM; KCl 25mM). (D) Fluorescence quantification of intracellular levels of K^+,^ Na^+^ and Ca^2+^ in hMDMs treated with ouabain and ouabain + KCl for 24 hours (ouabain 50 nM; KCl 25mM); n=3 donors. (E) Relative caspase 1 activity (left) and secreted IL-1β protein levels (right) in hMDMs treated with Ouabain (50 nM, 100 nM and, 500 nM) for 24 hours; at least n= 6 donors. (F) Relative caspase 3/7 activity (left) and secreted TNF*α* protein levels (right) in hMDMs treated with Ouabain (50 nM, 100 nM and 500 nM) for 24 hours; n=6 donors. (G) Cell viability (percentage of Annexin V^-^, PI^-^ cells) in hMDMs treated with ouabain and ouabain + caspase inhibitor (Q-VD-Oph) for 24 hours. Ouabain, 50 nM, 100 nM and 500 nM; Q-VD-Oph 2, 20μM; n=5 donors. Statistical significance was calculated using a paired one-way ANOVA test followed by Dunnett’s multiple comparisons (A, B, D, E and F) and student’s t-test (G). For (C), results are representative of at least 3 independent experiments. Error bars represent SEM. Scale bar= 250μm

It has been previously reported that ouabain, which contains a L-rhamnose unit (Supplementary Figure 1C), has higher affinity to the Na^+^,K^+^-ATPase than its aglycone ouabagenin, which lacks the L-rhamose unit (Cornelius et al., 2013). Accordingly, ouabagenin was not cytotoxic when treated for 24 hours at 50 nM, 100nM and 500 nM in hMDM (Supplementary Figure 1C), suggesting that the cytotoxic effects of ouabain were caused by inhibition of Na^+^,K^+^-ATPase. To test for the latter, we treated hMDMs with either ouabain (50nM) or ouabain in presence of KCl for 24 hours (Figure 1B). The usage of KCl rescued ouabain-induced cytotoxicity in hMDMs, confirming that cell death was caused by Na^+^,K^+^- ATPase inhibition and subsequent perturbation of intracellular ion balance in hMDMs. This was further investigated by incubating hMDMs treated with either ouabain alone or with KCl together with K^+^, Na^+^, Ca^2+^ fluorescent probes (Figure 1C). Intracellular K^+^ levels decreased in ouabain-treated cells compared to control hMDMS and intracellular K^+^ levels were partially rescued when KCl was added (Figure 1D). Similarly, intracellular levels of Na^+^ and Ca^2+^ increased in hMDMS treated with ouabain, which were also fully rescued by adding KCl (Figure 1D).

Calcium is a well-known mediator of the inflammasome activation in macrophages (Di et al., 2018). Ouabain has been recently reported to induce cell death by inflammasome and Caspase-1 activation in THP1 macrophage-like cells (LaRock et al., 2019). We thus evaluated the inflammasome activity by measuring luminescence resulting from catalytically active caspase-1 in the culture media of hMDMs. Caspase-1 activity was increased by 5-fold in ouabain-treated (50nM) compared to control hMDMs (Figure 1E). We next measured the levels of mature IL1*β* in culture media by ELISA and found a dose-dependent increase of IL1*β* in ouabain-treated hMDMs compared to control ones (Figure 1E). Ouabain also increased the activity of Caspase 3/7 and the release of the inflammatory cytokine TNF*α* in hMDM (Figure 1F). These results suggest that ouabain-induced cell death in hMDMs is associated with inflammatory and apoptotic caspases. To determine whether ouabain-induced cell death in hMDMs is caspase-dependent, hMDM were co-treated with ouabain and pan-caspase inhibitor Q-VD-OPh (Figure 1G). Caspase inhibition blunted the cytotoxic effect of ouabain, confirming a caspase-dependent cell death in ouabain-treated hMDMs.

### Ouabain depletes macrophages in the white adipose tissue ex vivo

To determine whether CGs induce cell death in tissue macrophages, human WAT from obese patients was cultured and treated with ouabain at nanomolar concentrations (50nM, 100nM, and 500nM) for 48 hours *ex vivo*. Macrophage-specific CD68 immunostaining showed a reduction of macrophages in ouabain-treated compared to control WAT explants (Figure 2A). This was confirmed by a ∼ 70% downregulation of *CD68* mRNA expression levels, measured by qRT-PCR, in ouabain-treated compared to control WAT explants (Figure 2B). Moreover, protein levels of the macrophage mannose receptor CD206 were assessed by Western Blot and found to be consistently reduced in ouabain-treated WAT explants compared to control ones (Figure 2C).

**Figure 2.**
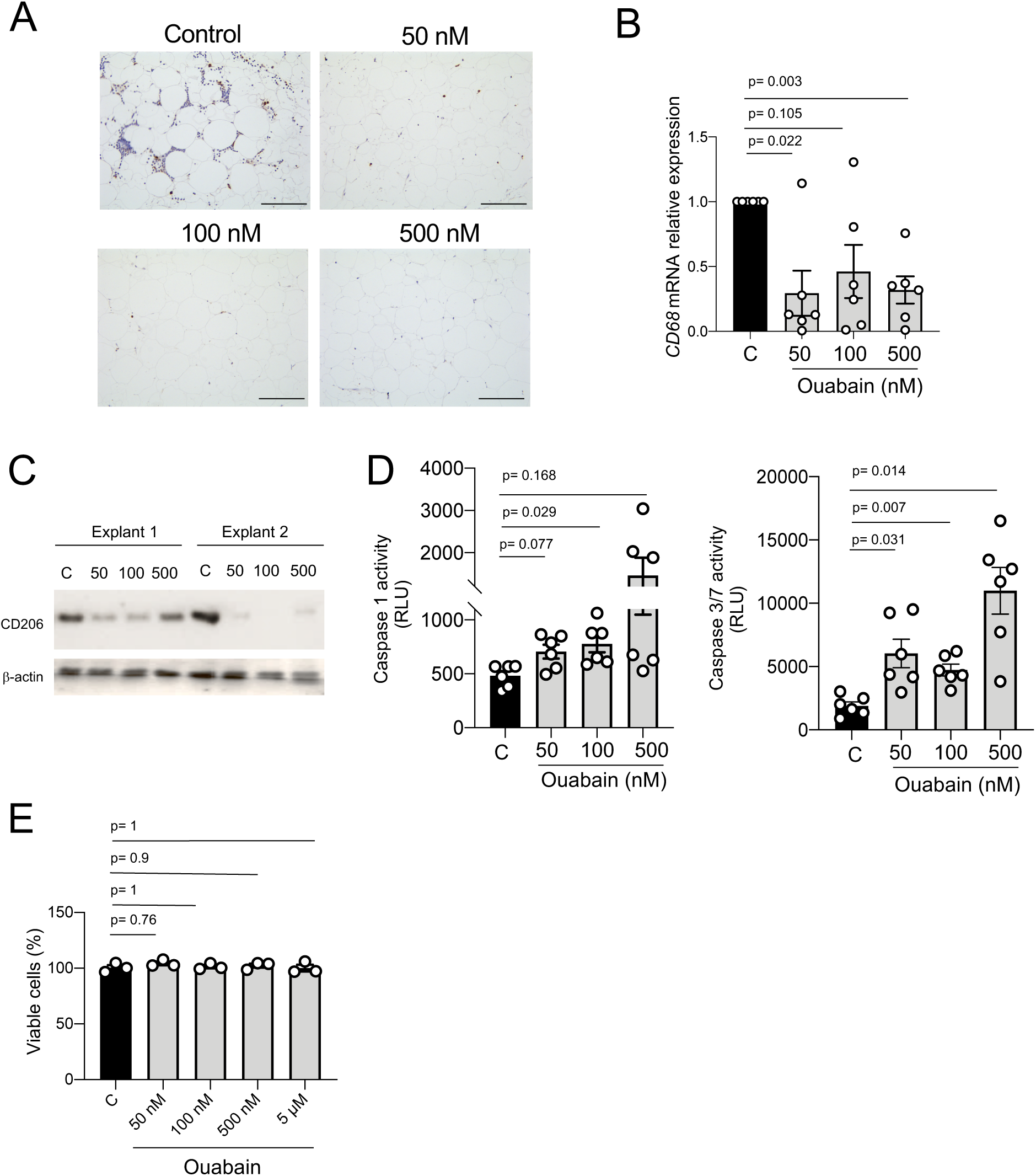
Ouabain depletes macrophages in human omental white adipose tissue (oWAT) explants. (A) Representative pictures of CD68 immunostaining in human oWAT explants treated with ouabain (50 nM, 100 nM and, 500 nM) for 48 hours. (B) Relative *CD68* mRNA expression in human oWAT explants treated with ouabain (50 nM, 100 nM and 500 nM) for 48 hours, n=6 explants. (C) CD206 Western blotting in human oWAT explants treated with ouabain (50 nM, 100 nM and, 500 nM) for 48 hours. (D) Relative caspase 1 (left) and caspase 3/7 (right) activity in human oWAT explants treated with ouabain (50 nM, 100 nM and, 500 nM) for 48 hours; at least n=6 explants (E) Cell viability (AlamarBlue reduction) of SVF-derived adipocytes treated with ouabain (50 nM, 100 nM and, 500 nM) for 48 hours. Statistical significance was calculated using paired one-way ANOVA test followed by Dunnett’s multiple comparisons. Error bars represent SEM. Scale bar= 100 μm

To determine whether the ouabain-induced *ex vivo* macrophage depletion was due to the activation of inflammatory and apoptotic caspases, (as previously seen in hMDMs – Figure 1), the activity of caspases 1 and 3/7 was measured in WAT explants culture media (Figure 2D). Caspases 1 and 3/7 activity was higher in ouabain-treated compared to control WAT explants, suggesting that ouabain induced macrophage cell death through pyroptosis and apoptosis in WAT explants.

In order to evaluate the potential cytotoxicity of ouabain in non-immune resident cells such as adipocytes, human WAT-derived stromal cells were differentiated into adipocytes and treated with ouabain (50nM, 100nM,500nM and 5μM) for 24 hours (Figure 2E). Ouabain cytotoxicity was measured by Alamar blue reduction assay and showed no differences among all the groups, indicating that ouabain had no impact on adipocyte viability.

### Ouabain improves insulin sensitivity and mediates ECM remodeling in the WAT ex vivo

Macrophage infiltration is a hallmark of inflammation and insulin resistance in visceral WAT depots in mice and humans (Koppaka et al., 2013; Olona et al., 2018; Wentworth et al., 2010). Insulin sensitivity was assessed by measuring the levels of AKTser473 phosphorylation, a major effector of insulin signaling, in control and ouabain-treated WAT explants (Figure 3A). Ouabain treatment increased AKTser473 phosphorylation by ∼50% compared to control WAT explants (Figure 3B), indicating that ouabain promotes insulin sensitivity in the *ex vivo* cultured WAT. We then confirmed this finding by measuring *ADIPOQ* mRNA expression and CD36 protein levels by qRT-PCR and Western blot, respectively. *ADIPOQ* encodes for Adiponectin, an adipokine that promotes insulin sensitivity (Shetty et al., 2009), and was significantly up-regulated in ouabain-treated WAT explants compared to control ones (Figure 3C). Similarly, protein levels of CD36, which have an important role in fatty acid uptake upon insulin stimulation, was also increased in ouabain treated WAT explants (Figure 3D). We next investigated whether increased CD36 and fatty acid uptake could affect adipocyte size, as previously reported (Vroegrijk et al., 2013). Adipocyte mean areas were ∼20% bigger in ouabain-treated compared to control WAT explants (Figure 3E). These results show that ouabain improves tissue function, promoting insulin sensitivity and fatty acid uptake in WAT from obese patients.

**Figure 3.**
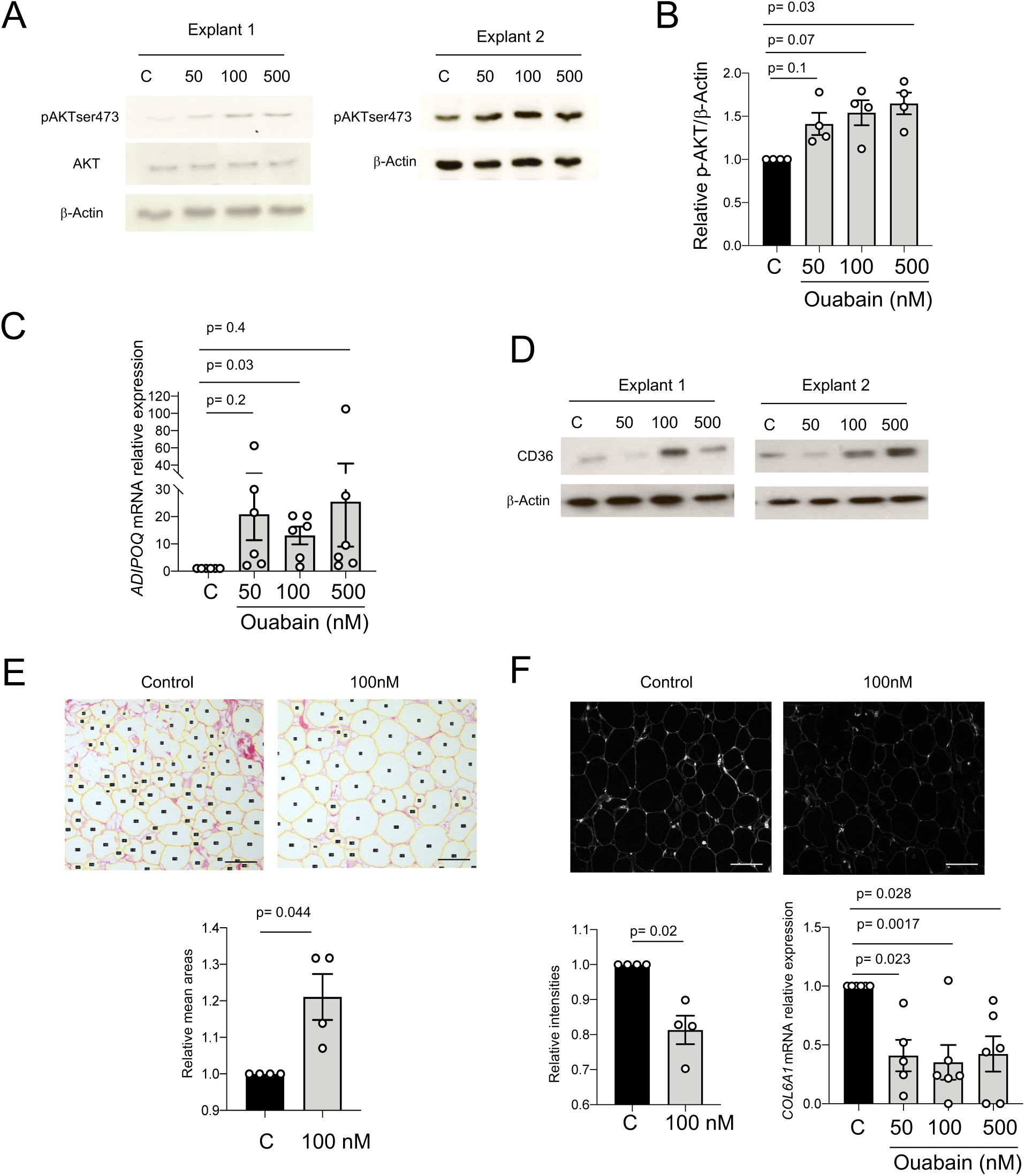
Ouabain improves insulin sensitivity and reduces Type VI collagen deposition in human omental white adipose tissue (oWAT) explants. (A) Phospho-AKTser473, total AKT, and β-actin Western blotting in human oWAT explants treated with insulin and ouabain (50 nM, 100 nM and, 500 nM) for 48 hours. (B) Relative levels of phospho-AKTser473 protein, normalized to the loading control (β -actin); n=4 explants. (C) Relative *ADIPOQ* mRNA expression in human oWAT explants treated with ouabain (50 nM, 100 nM and, 500 nM) for 48 hours; n=6 explants. (D) CD36 Western blotting in human oWAT explants treated with ouabain (50 nM, 100 nM and, 500 nM) for 48 hours. (E) Representative Sirius Red pictures of human oWAT treated with ouabain (100 nM) for 48 hours (top). Relative mean adipocyte areas in human oWAT explants treated with vehicle (control) and ouabain (100 nM) for 48 hours (bottom); n=6 explants. (F) Representative type VI collagen immunofluorescence pictures of human oWAT treated with ouabain (100 nM) for 48 hours (top). Relative type VI collagen fluorescence (bottom left) and relative *COL6A1* mRNA expression in human oWAT explants treated with vehicle (control) or ouabain (50 nM, 100 nM and, 500 nM) for 48 hours; at least n=4 explants. Statistical significance was calculated using paired one-way ANOVA test followed by Dunnett’s multiple comparisons (B, C, F) and one sample t-test (E and F). Error bars represent SEM. Scale bar= 50 μm

Adipose tissue macrophages have been shown to regulate collagen deposition and fibrosis in visceral WAT during obesity, inducing tissue remodelling and insulin resistance (Keophiphath et al., 2009). In line with this, type VI collagen knockout mice upon high-fat diet showed unrestrained adipocyte expansion and improved WAT insulin sensitivity (Khan et al., 2009). Type VI collagen immunostaining was performed and mean grey intensities showed a down-regulation of type VI collagen in ouabain-treated compared to control WAT explants (Figure 3F). This coincided with *COL6A1* mRNA expression downregulation (Figure 3F), confirming a decrease of type VI collagen synthesis and deposition in ouabain-treated compared to control WAT explants.

## Discussion

Here we show that nanomolar concentration of CGs causes caspase-1 mediated pyroptosis in human macrophages *in vitro*. In the white adipose tissue (WAT) explants isolated from obese patients with T2DM, the same treatment causes macrophage depletion. We also observe that CG-dependent macrophage cytotoxicity results in TNF secretion *in vitro*, which we could not detect in the ex vivo WAT (data not shown). Collectively, these results show that CG-mediated decrease in intracellular K^+^ and the net increase in Ca^2+^ levels trigger both pyroptotic (Caspase-1-dependent IL-1β secretion) and apoptotic (TNF-mediated cell death through caspase 3) cell death pathways in human macrophages.

Pyroptosis is caspase-1-dependent and causes the assembly of the inflammasome complex which initiates the efficient release of IL-1β (Gross et al., 2011). While pyroptosis is an inflammatory form of cell death triggered by caspase-1, apoptotic cell death is mediated by effector apoptotic caspases such as caspase-3, −6, and −7 (Man and Kanneganti, 2016). Recent evidence shows a significant crosstalk and compensation between apoptotic and pyroptotic cell death mechanisms and caspases such as caspase-6 having master-regulatory role in both events (Zheng et al., 2020). Hence it is likely that CGs trigger broader caspase activation pathways in human macrophages which causes IL-1β and TNF release, in line with reports showing TNF secretion following co-treatment with LPS and ATP (Barbera-Cremades et al., 2017; Di et al., 2018). Interestingly TNF secretion has been previously shown to be Caspase-1 dependent in macrophages (Miggin et al., 2007), which supports the broader effects of inflammasome activation in macrophages.

We report that the cell death caused by CGs in human macrophages is dependent on Na^+^,K^+^-ATPase activity and requires the presence of the sugar moiety attached to the steroid part of the compound. Indeed, the addition of KCl in the media showed an almost complete rescue of cell viability and intracellular levels of Na^+^ and Ca^2+^. This also confirms that, similar to cardiomyocytes, macrophage survival is tightly dependent on potassium efflux and the sensitivity to CGs is more prominent in human macrophage-like cells when compared to murine ones (LaRock et al., 2019). Furthermore, our data suggest that, among other peripheral blood mononuclear cells, CGs seem to have a particularly potent effect on macrophages. This leads to the question of *why human macrophages among other cells undergo apoptosis and pyroptosis as a result of a decrease intracellular potassium*? Interestingly, CGs such as digoxin and digitoxin selectively induce apoptosis of senescent cells and digitoxin have a senolytic (i.e. selective killing of senescent cells) activity at a nanomolar range concentration that is closed to the one observed in cardiac patients treated with this drug (Guerrero et al., 2019; Lopez-Lazaro, 2007). This raises the possibility of phenotypic similarities between senescent cells and macrophages and a recent report shows that chemotherapy-induced senescent breast cancer cells are highly enriched for macrophage genes and can perform phagocytosis (Tonnessen-Murray et al., 2019). It is therefore tempting to speculate that the macrophages and senescent cells show a shared sensitivity to the CG-mediated apoptosis, although the underlying mechanisms may differ (induction of pro-apoptotic Bcl2 in senescent cells vs. pyroptosis/apoptosis in macrophages).

When used at nanomolar concentrations, CGs showed effective depletion of macrophages from *ex vivo* cultured white adipose tissue explants isolated from obese patients undergoing bariatric surgery. We observed an almost complete down-regulation of CD206 (mannose receptor). Indeed, adipose tissue-infiltrating macrophages are responsible for a persistent low-grade inflammatory state that underlies the systemic insulin resistance observed in obesity and CD206 is a specific marker for adipose tissue macrophages which controls adipogenesis (Nawaz et al., 2017). In fact, the partial depletion of CD206+ macrophages enhances insulin sensitivity (Igarashi et al., 2018) and we show that this is indeed the case with the usage of ouabain (50nM-100nM) in the *ex vivo* cultured WAT explants. This WAT macrophage depletion could be partly through Caspase 1-mediated pyroptosis, though we cannot exclude apoptotic cell death despite no increase in tissue TNF levels.

We report an improved WAT function upon treatment with ouabain. Inflammation and fibrosis are two major pathways that dysregulate WAT homeostasis in obesity, and current therapeutic strategies aim to target these pathways (Kusminski et al., 2016). Our findings show that ouabain-mediated macrophage cytotoxicity increases adipocyte hypertrophy and induces metabolic activity through up-regulation of CD36, which facilitates the uptake of long-chain fatty acids. (Christiaens et al., 2012). Moreover, ouabain treatment reduces the type VI collagen, the main collagen in the WAT which accumulates following metabolically challenging conditions (Khan et al., 2009). Interestingly, macrophages have been shown to produce type VI collagen in the lung (Ucero et al., 2019), which suggests that ouabain-mediated macrophage depletion is the direct cause of the decreased levels of this collagen in the WAT. We show that SVF-derived cultured adipocytes are not sensitive to ouabain-induced cell death and our data obtained in the peripheral blood suggests that macrophages are likely to be the only resident immune cells in the WAT that are cleared from the tissue through mechanisms that remain to be identified. Furthermore, the improvement of WAT homeostasis following *ex vivo* treatment with ouabain can be through soluble factors secreted by dying macrophages and/or direct effects on adipocytes. For instance, adipocytes treated with ouabain showed a possible non-canonical role of this CG through up-regulation of Glut4 in these cells (Brewer et al., 2019). The identification of all soluble factors accompanying ouabain-mediated macrophage death and elucidating the pathways triggered by CGs in adipocytes will be important in fully dissecting the mechanisms involved in the beneficial role of these compounds in inflamed tissues.

Here we show the beneficial role of the usage of ouabain in a tissue where the increase in macrophage infiltrates is associated with tissue damage and inflammation. In an infectious disease context, the use of CGs can be detrimental because of the bactericidal role of mononuclear phagocytes (Esposito, 1985). Hence the therapeutic advantage of nanomolar range usage of CGs can be studied in sterile inflammation where tissue macrophage presence is generally associated with poor clinical outcome (e.g. inflammatory brain disorders, autoimmune disease). In keeping with this, the local administration of CGs can have therapeutic effects through selective cytotoxicity towards tissue-resident macrophages, bypassing the unwanted side effects related to the reported toxicity of these compounds. Considering that some CGs such as digoxin are frequently prescribed medicines in elderly population, these findings will also re-evaluate the usage of these in the context of tissue macrophage integrity in health and disease.

---

**Supplementary Figure 1.**
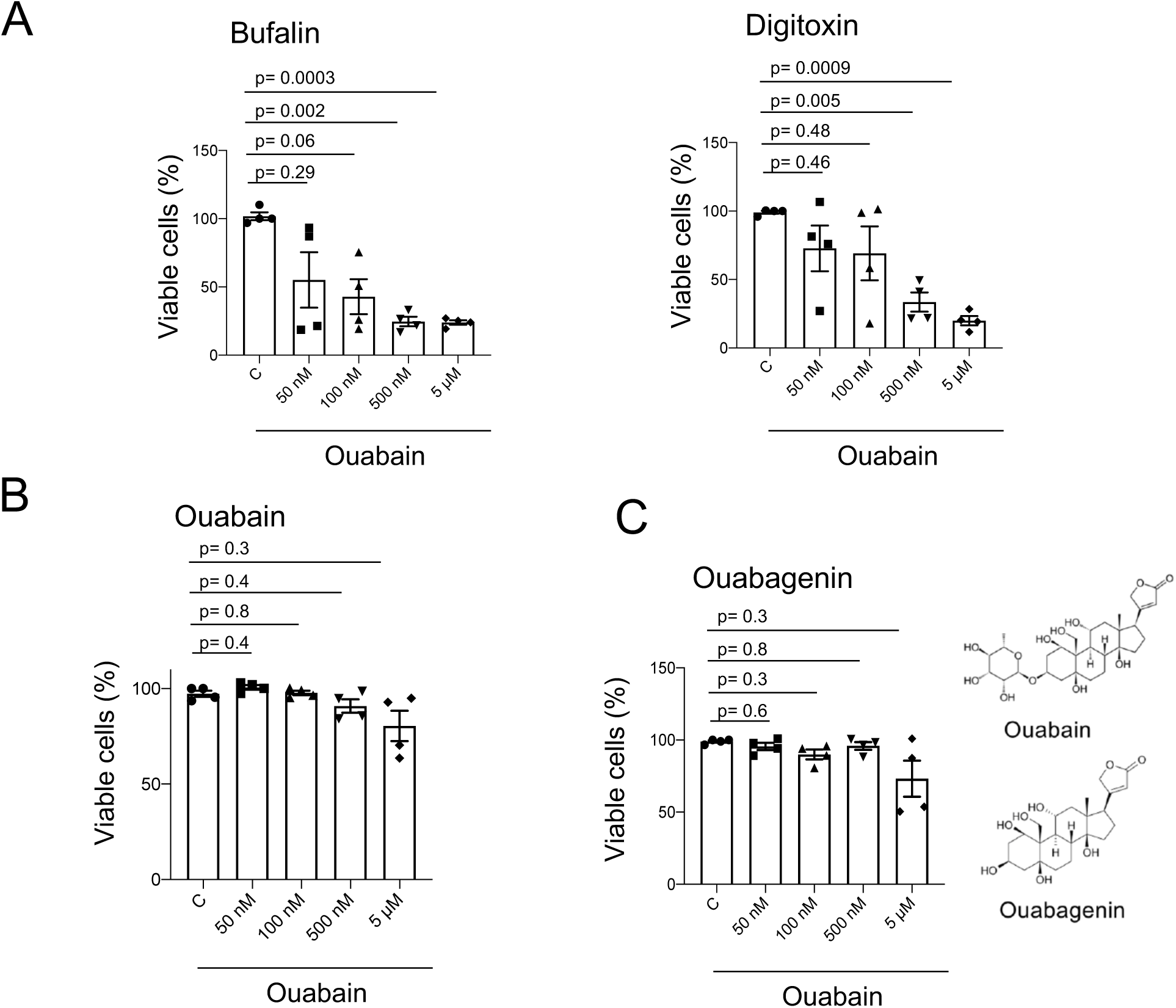
Cardiac glycosides induce caspase-dependent cell death by inhibiting the Na^+^,K^+^-ATPase in human monocyte-derived macrophages (hMDMs). (A) Cell viability (percentage of Annexin V^-^,PI^-^ cells) in hMDMs treated with bufalin (left) and digitoxin (right) at 50 nM, 100nM, 500 nM or 5 μM for 24 hours; n=4 donors. (B) Cell viability (percentage of Annexin V^-^,PI^-^ cells) in non-adherent PBMCs treated with ouabain at (50 nM, 100nM, 500 nM and 5 μM) for 24 hours; n=4 donors. (C) Cell viability (percentage of Annexin V^-^,PI^-^ cells) in hMDMs treated with ouabagenin (50 nM, 100nM, 500 nM and 5 μM) for 24 hours; n=4 donors. Statistical significance was calculated using paired one-way ANOVA test followed by Dunnett’s multiple comparisons. Error bars represent SEM.

